# High throughput genome-wide single cell protein:DNA binding site mapping by targeted insertion of promoters (TIP-seq)

**DOI:** 10.1101/2021.03.17.435909

**Authors:** Daniel A. Bartlett, Vishnu Dileep, Steve Henikoff, David M. Gilbert

## Abstract

Assessing cell to cell, and importantly, chromosome to chromosome, heterogeneity in cellular phenotypes is a central goal of modern cell biology. However, chromatin profiling in single cells has been extremely challenging, and single chromosome profiling has not been achieved. In cases where single cell methods have shown promise, success has been mainly limited to histone proteins and/or require highly specialized equipment or cell type specific protocols and are relatively low throughput. Here, we have combined the advantages of tagmentation, linear amplification and combinatorial indexing to produce a high throughput single cell DNA binding site mapping method that does not require specialized equipment and is capable of multiplexing several samples/ target proteins in one experiment. Targeted Insertion of Promoters (TIP-seq) uses Tn5 fused to protein A (as with CUT&Tag) to insert a T7 RNA polymerase promoter into sites adjacent to an antibody bound to a chromatin protein of interest, followed by linear amplification of flanking DNA with T7 polymerase, cDNA preparation and PCR indexing. Tip-seq provides ∼10-fold higher unique reads and thus higher coverage per single cell compared to state-of-the-art methods. We apply TIP-seq to map histone modifications, RNA PolII and CTCF binding sites in single human and mouse cells. TIP-seq will also be adaptable for other platforms, such as 10X genomics and ICELL8. In summary, TIP-seq provides a high-throughput, low-cost method for single cell protein mapping, that yields substantially higher coverage per cell and signal to noise than existing methods.

## Introduction

The surge in single cell epigenetics and transcriptomics has been instrumental to our growing understanding of cell fate changes and the nature of cell heterogeneity during development and disease. Unfortunately, tracing the role of chromatin-protein binding in single-cell context remains uncharted territory due to the lack of efficient and robust methods to map the binding sites of chromatin proteins in single cells. The mounting demand for a highly efficient, robust, and scalable method to map protein binding sites genome-wide in low- and single-cells is evident in the massive proliferation of low input ChIP methods (Brind’Amour et al., 2015; Cao et al., 2015; van Galen et al., 2016; Grosselin et al., 2019; Zhang et al., 2016), but ChIP methods all suffer from the inherent inefficiency of immunoprecipitation (IP) (Baranello et al., 2016; Marinov, 2018). DamID identifies binding sites via transgenic expression of DNA methyltransferase fused to the target-protein of interest that methylates nearby adenines, and then subsequent cutting of DNA fragments specifically at DpnII restriction enzyme cut sites. DamID bypasses the inefficiency of IP, but requires cloning, transfection, and in situ expression of the transgenic Dam-fusion protein, its resolution is limited by the distribution of DpnII cutting sites and it requires enzymatic end-prep and adapter ligation steps prior to library amplification that lead to sample loss. Methods such as CUT&RUN (Hainer et al., 2019; Skene et al., 2018) and scChIC-seq (Ku et al., 2019) tether micrococcal nuclease proteins to the target protein of interest to cleave and solubilize the surrounding chromatin. These approaches also avoid IP, but still require library prep steps that result in sample loss. Tagmentation via Tn5 transposase simultaneously cleaves DNA and attaches Illumina sequencing adapters in a one-step *cut-and-paste* mechanism. Methods such as CUT&Tag (Kaya-Okur et al., 2019), CoBATCH (Wang et al., 2019) and ACT-seq (Carter et al., 2020) recruit a proteinA-Tn5 fusion protein to the antibody-bound target protein of interest to produce libraries of fragments that can be subjected to PCR amplification.

A recent chromatin accessibility profiling method demonstrated that modifying the Tn5 transposon to deliver a T7 promoter in place of standard Illumina PCR adapters enabled >1000 fold linear amplification of adjacent gDNA by T7 RNA polymerase prior to reverse transcription and cDNA library preparation (Sos et al., 2016). Linear amplification prior to PCR library amplification reduced input requirements for profiling DNaseI hypersensitive sites from 50,000 down to single cells (Lake et al., 2018). Linear amplification also provides unbiased genomic coverage since only a single ligation event is required to amplify any given segment (Chen et al., 2017; Rooijers et al.), whereas PCR requires two ligation events to occur within an optimal distance and in the correct orientation to enable PCR amplification. Linear amplification has been integrated with other methods and shown to significantly increase library DNA yield and detection sensitivity when substituted for conventional amplification (Chen et al., 2017; van Galen et al., 2016; Harada et al., 2019; Hashimshony et al., 2012; Lake et al., 2018; Rooijers et al.). For example, Chromatin Integration Labeling (ChIL) uses a secondary antibody tethered to an oligo containing a T7 promoter upstream of the Tn5 mosaic binding sequence, and in-situ tagmentation inserts the T7 promoter into adjacent DNA. ChIL provides impressive sensitivity, and has mapped protein-binding sites of select histone modifications in single mouse cells, and TF CTCF in 100 cells (Harada et al., 2019). Unfortunately, ChIL is only compatible with stongly adherent cell-culture samples, is not amenable to robotic automation and requires in-house construction of antibody-oligo conjugates.

We wished to develop a single cell chromatin profiling method that could be applied to any sample type and scaled to high throughput. CUT&Tag is compatible with any sample-type, is readily automatable, fixation independent, inexpensive and simple to perform and has shown some promise with single cell profiling (Kaya-Okur et al., 2019). Our method, <u>***T**argeted **I**nserton of **P**romoters (Tip-seq)*</u>, inserts T7-promoters adjacent to all antibody-bound genomic sites to permit subsequent linear amplification of adjacent genomic DNA. The use of *in vitro* transcription in TIP-seq leads to substantially increased sensitivity over standard PCR library preparation employed in CUT&Tag. For single cell mapping, the primary obstacle with all state-of-the-art methods has been sparse genome coverage (Minkina and Shendure, 2019). TIP-seq overcomes this obstacle with a remarkable (10-fold) improvement in library complexity per single cell, achieving high signal to noise single cell data. To increase the throughput of TIP-seq and avoid the expense and equipment-dependency of single-cell platforms, we further adapted the method to combinatorial Indexing (sciTIP-seq), which will allow high throughput (theoretically up to 38,500 per run) and the ability to profile multiple samples and antibody targets. In summary, sciTIP-seq provides a high throughput, low-cost, low background method for single cell protein mapping with substantial gains in terms of per-cell read coverage.

## RESULTS

### TIP-seq design

The excellent sensitivity of CUT&Tag is owed to its use of Tn5 to streamline library preparation by directly inserting PCR sequencing adapters in-situ, but paradoxically its sensitivity is inherently limited by PCR. Since Tn5 inserts adapters (ME-A/B) in random orientations, roughly half the targets do not have adapters in the correct orientation to amplify. Further, PCR library prep is extremely sensitive to size variations of amplicons: When two adjacent transposition events occur too far apart, they will not amplify efficiently during PCR or sequencing cluster generation, but if too close, they will exponentially bias library coverage due to increased PCR amplification and clustering efficiency of short fragments. Target regions that are small in length (ie. TF binding sites) also contend with having a decreased likelihood of receiving multiple transposition events to overcome the 50% efficiency due to adapter orientation. Linear amplification has long been used to circumvent amplification and sequence composition bias resulting from PCR amplification and is particularly beneficial when limited starting DNA is available (van Bakel et al., 2008; Eberwine et al., 1992; Hoeijmakers et al., 2011). In fact, numerous chromatin-mapping methods have transitioned to linear amplification for increased sensitivity in single-cells including scRNA-seq (a.k.a. CEL-seq; Hashimshony et al., 2012), ChIL-seq (Harada et al., 2019), MINT-ChIP (van Galen et al., 2016), DamID (Rooijers et al.), ATAC-seq (a.k.a THS-seq; Lake et al., 2018), and whole genome sequencing (a.k.a LIANTI; Chen et al., 2017). Therefore, we hypothesized that adapting CUT&Tag to use linear amplification would greatly improve the sensitivity in single cells. By merging CUT&Tag with existing Tn5 transposition-based linear-amplification protocols, we created a custom-designed proteinA-Tn5 transposase fusion protein (pA-Tn5), that carries with it a transposon containing a T7 promoter (**Figure 1B**; see supplementary data for custom oligos; adapted from (Harada et al., 2019; Lake et al., 2018), that is recruited to antibody-bound chromatin sites where the Tn5 transposome simultaneously cuts and inserts (tagmentation) the T7 promoter into adjacent genomic DNA to permit linear amplification (**Figure 1**). In-vitro transcription (IVT) by T7 RNA polymerase creates roughly 1000-fold RNA copies of insertion sites, and after reverse transcription, second-strand synthesis, and cDNA fragmentation, TIP-seq amplicons are prepared for sequencing via limited-cycle number PCR indexing. With TIP-seq, the distance between two transposition sites is irrelevant since only one T7 promoter insertion is required to amplify the site. We also adapted the pA-Tn5 transposon to deliver only T7-promoter containing adapters (rather than Forward & Reverse PCR adapters) thus completely mitigating the efficiency loss of mismatched PCR adapter orientations and distance between insertions. Additionally, linear amplification produces higher fidelity and uniformity since mistakes made during amplification do not themselves become templates to exponentially propagate the mistakes, which translates to higher mappability of single-cell sequencing reads (Chen et al., 2017). Furthermore, TIP-seq is a single-tube protocol to reduce potential sample loss.

**Figure 1.**
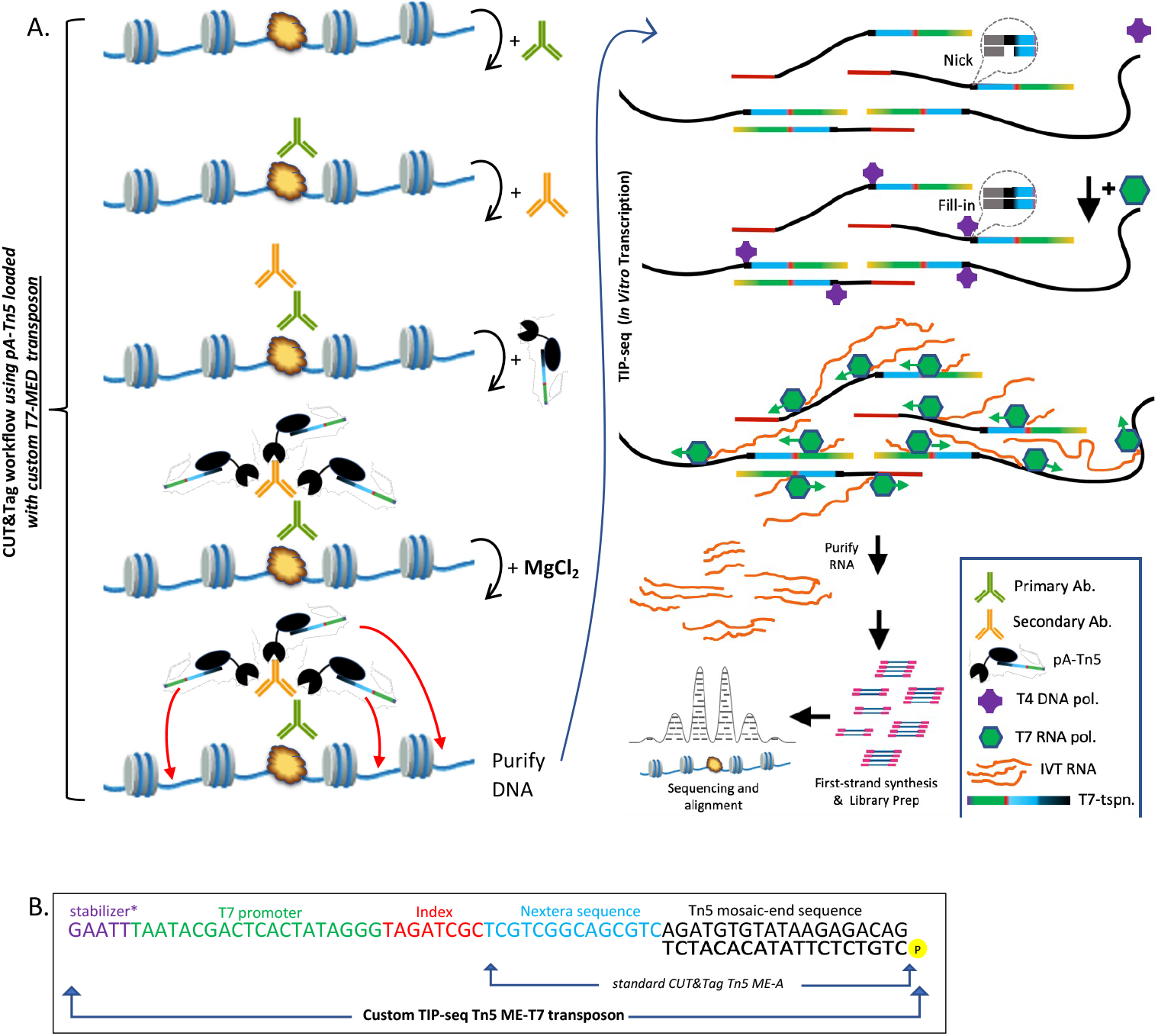
Overview of Targeted Insertion of Promoters (Tip-seq) A robust low-cell A robust low-cell mapping method that combines CUT&Tag with RNA-mediated linear amplification. (A) See text for details. (B) Custom T7-MED transposon used in TIP-seq to insert T7 promoters near antibody bound targets. Standard CUT&Tag MED-A shown for comparison. *upstream AT-rich stabilizer sequence increases affinity of T7 RNA Polymerase for promoter and the efficiency of promoter clearance (Tang et al. 2005).

### Linear amplification substantially increases sensitivity and library coverage over conventional PCR library preparation

To assess the advantage of linear amplification over standard PCR library preparation we performed TIP-seq and CUT&Tag in parallel on decreasing numbers of cells for histone modification H3K27me3 and the transcription factor CTCF in human HCT116 cells (**Figure 2A-B**). Enhanced efficiency was already evident by a ∼183-fold average increased yield of CTCF library DNA for TIP-seq vs. CUT&Tag. Library prep and sequencing of CTCF TIP-seq samples produced excellent results for all cell numbers attempted, whereas CUT&Tag libraries failed to yield sufficient library DNA for sequencing in 1000 or fewer cells, and sequencing reads from CTCF TIP-seq using 5000 cells yielded ∼45-fold more unique (non-duplicated) fragments compared to their CUT&Tag counterparts (**Figure 2C)** indicative of increased sensitivity and efficiency. We then compared the enrichment of reads for the CTCF samples surrounding all CTCF peaks (+/−2kb) called using Bulk ChIP-seq (ENCODE). Compared to 5000 cell CUT&Tag, the enrichment of reads at ENCODE CTCF peaks was much higher in 5000 cell (∼1.8-fold) and 1000 cell (1.3-fold) TIP-seq (∼1.8-fold and ∼1.3 fold, respectively) and a higher proportion of the ChIP peaks displayed read occupancy (**Figure 2D**). Remarkably, 100 and 50 cell TIP-seq peak enrichment was comparable to 5000 cell CUT&Tag.

**Figure 2.**
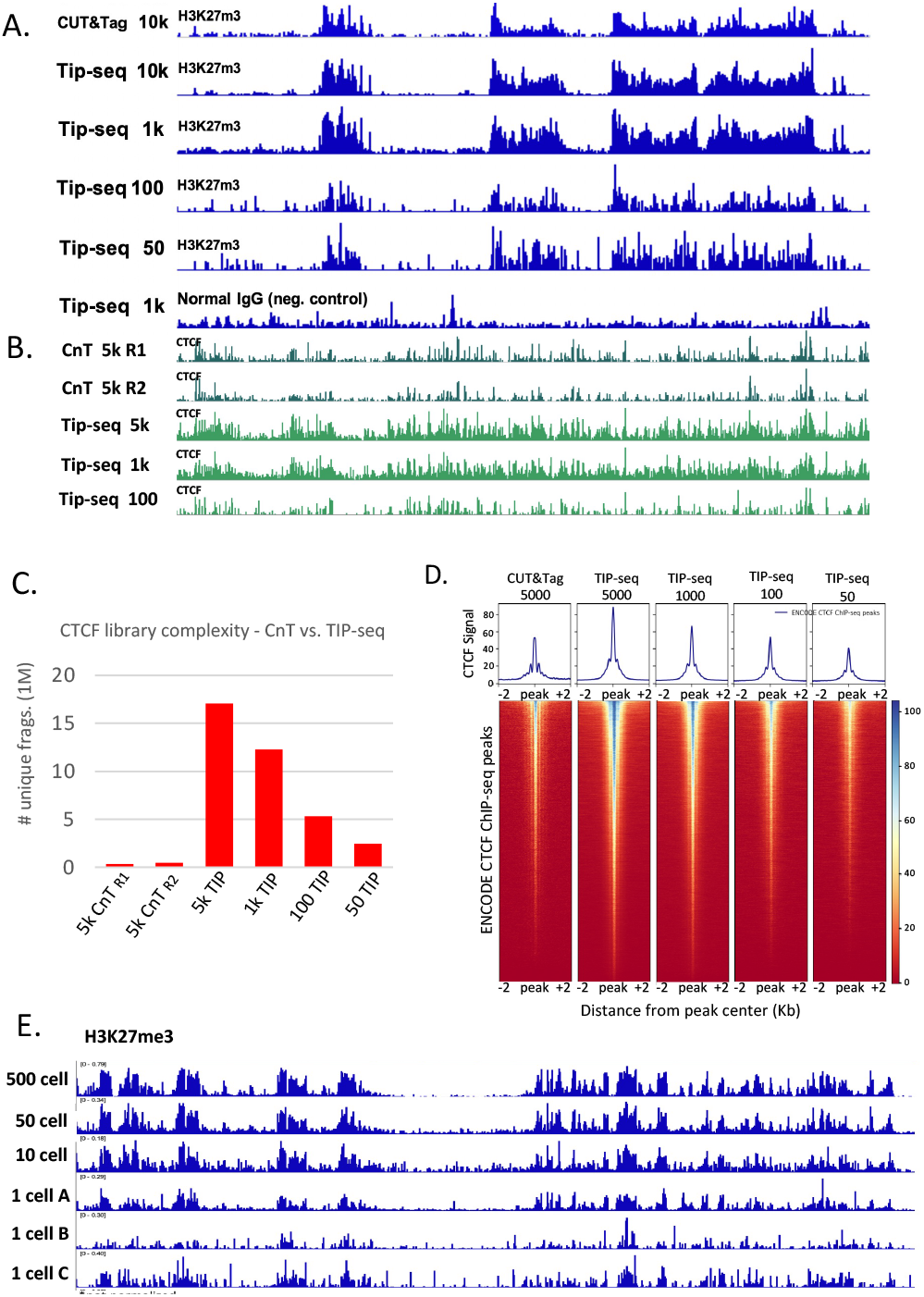
Bulk TIP-seq substantially increases library complexity and sensitivity over PCR based library prep. (A) **Histone modifications**. IGV track view across a 3 Mb segment of the human genome for H3K27me3 TIP-seq on 10k, 1k, 100, and 50 HCT116 cells. Tracks show CUT&Tag data in 10k cells for comparison. TIP-seq for Normal IgG in 1k cells shown as negative control. (B) **Transcription factors**. IGV track view across a 3 Mb segment of the human genome for CTCF TIP-seq in 5k, 1k, and 100 HCT116 cells. Tracks show CUT&Tag data in 5k cells for comparison. (C) **Library complexity of CTCF TIP-seq vs CUT&Tag** as a fraction of unique (non-duplicated) reads over total reads. Samples were processed in parallel, with CUT&Tag pA-Tn5 loaded with ME-A/B adapters, or TIP-seq pA-Tn5 loaded with ME-T7 transposons, and following Tagmentation and DNA purification, CUT&Tag DNA was PCR amplified using 15 cycles, while TIP-seq DNA was processed as described (Fig1) and indexed with 9 PCR cycles. Libraries were pooled to equimolar ratios and paired-end sequenced. (D) **Enrichment heatmaps** of CTCF TIP-seq vs CUT&Tag surrounding +/−2kb ENCODE CTCF peaks. Samples were normalized to the sum of per-base read coverage and scaled to 1x genome coverage before plotting heatmaps with deepTools. (E) **Single cells**. IGV track view showing TIP-seq data collected from single cells. Cells were processed in bulk until terminating tagmentation reaction, then cells were FACS sorted into individual PCR tubes for DNA purification and subsequent in-vitro transcription, cDNA library prep, and sequencing.

### Improved read coverage of TIP-seq permits robust mapping of epigenetic marks in single cells

All efficiency losses and bias accumulation caused by PCR are exacerbated when limited input material is used. Therefore, we next sought to test how well TIP-seq worked with a single cell input. To do this, we performed TIP-seq in bulk (but omitted the use of magnetic conA magnetic beads) up until termination of the tagmentation reaction. Next, we FACS sorted single cells into PCR tube strips and proceeded as normal with the TIP-seq protocol. Three of the five single cells yielded ample DNA to be pooled for sequencing. Again, those three cells delivered profiles remarkably resembling those of bulk samples, albeit with varying signal and background (**Fig 1E**).

### High-throughput, low-cost single-cell combinatorial indexing TIP-seq (sciTIP-seq)

A limitation of conventional single-cell preparatory methods is that single cells must be physically separated and compartmentalized before being biochemically processed in separate reaction volumes causing cost and labor intensity to prohibitively scale linearly with the number of single cells processed. Furthermore, working with single cells, small volumes and low DNA inputs leads to sample loss. These problems are somewhat mitigated by using specialized microfluidic handlers (Refs.) but availability and access to expensive and specialized equipment limits the adoption of such techniques. By contrast, single-cell combinatorial indexing (‘sci’) is a highly scalable method that that has been widely adopted to acquire profiles of transcriptomes, genomes, chromatin accessibility, methylomes and chromosome conformation in tens to hundreds-of-thousands of single cells without the need for compartmentalization of individual cells (Cao et al.; Cusanovich et al., 2015; Lareau et al., 2019; Mulqueen et al., 2018; Ramani et al., 2017; Vitak et al., 2017; Wang et al., 2019; Weiner et al., 2016; Zhu et al., 2021). Throughput is high and cost-per-cell decreases exponentially with the number of cells processed. Sci workflows thus permit affordable and highly scalable throughput, require no specialized equipment that is not widely available, and sci workflows inherently enable sample multiplexing.

We adapted TIP-seq for combinatorial indexing based on existing tagmentation-based sci-protocols (Cusanovich et al., 2015; Lake et al., 2018; Lareau et al., 2019) (**Figure 3**). Briefly, cells were incubated with primary and secondary antibodies in bulk before being split into a 96 well plate (∼2000 cells/well) for the first round of indexing. To add the first index, we created 384 uniquely-barcoded T7-MEDS transposons (adapted from (Lake et al., 2018); T7-promoter optimized based on (Tang et al., 2005)) such that cells in each well of a 96 well plate receives a unique index downstream of the T7 promoter (up to 4 plates can be processed for Index1-384 MEDS). After addition of Index 1 via tagmentation with the custom pA-Tn5 transposons, cells were pooled and redistributed randomly via FACS into 96 well plates (15-100 cells/well). DNA was purified and subjected to IVT and cDNA prep as described for bulk TIP-seq. Index 2 was incorporated via PCR resulting in library fragments (**Figure 3B**) containing both an r5 barcode (added via pA-Tn5) and an i5 and i7 barcode (added via PCR). The resulting libraries were pooled to equimolar ratios and paired-end sequencing was performed with 29 cycles for index i5 (8 cycle index run by default) to capture all single-cell index sequences that permit retroactive demultiplexing of reads back to their individual cells of origin.

**Figure 3.**
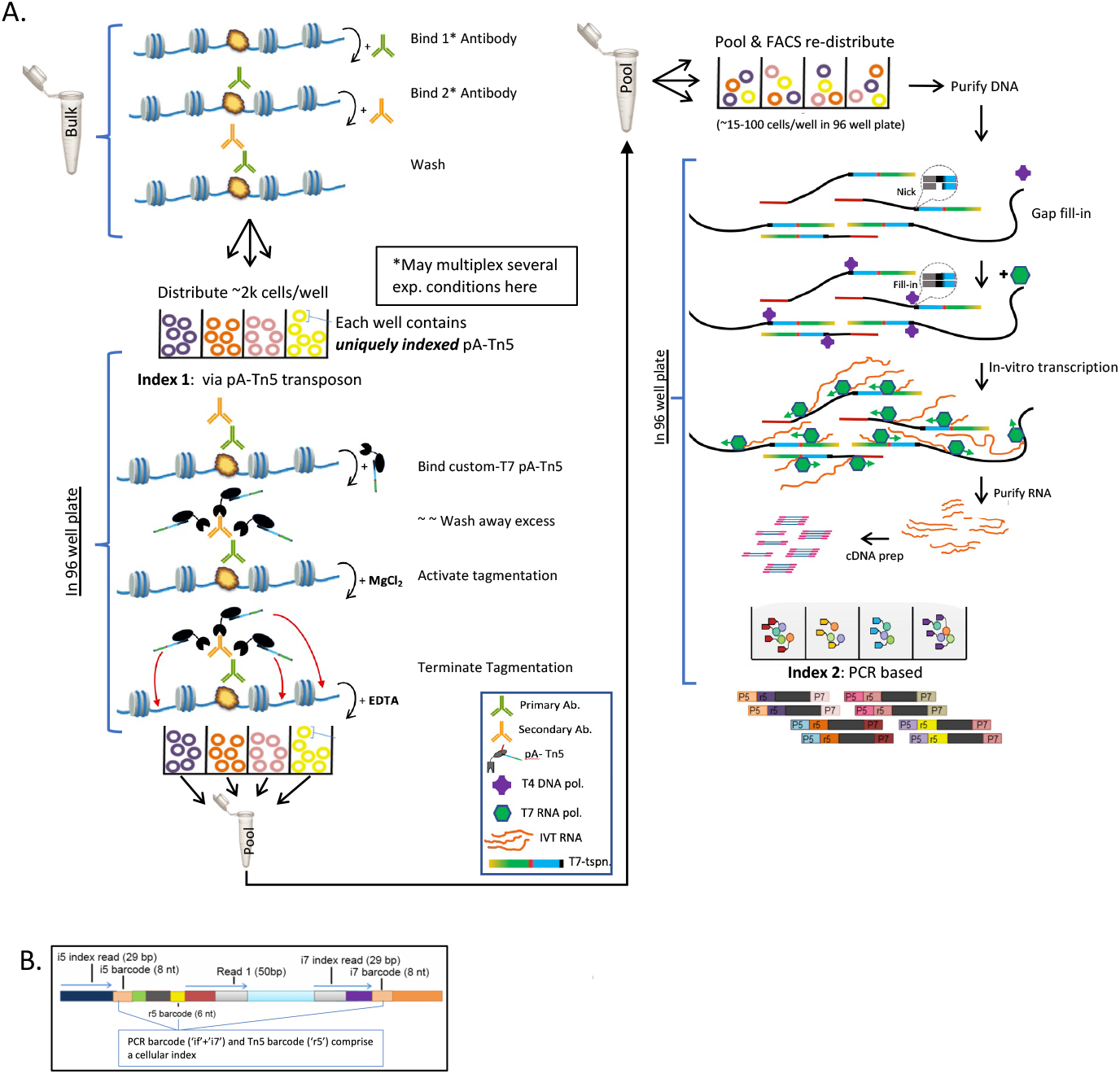
Overview of combinatorial indexing TIP-seq (sciTIP-seq). (A) Cells are harvested, permeabilized, and treated with primary and secondary antibodies in bulk. Cells are counted and distributed to a 96 well plate (∼2000 cells per well) where they are incubated with custom uniquely indexed pA-Tn5. Cells are washed to remove unbound pA-Tn5 before activating tagmentation. Tagmentation is terminated by addition of EDTA, and cells are pooled together and redistributed at random to new 96 well plate (∼15-100 cells per well *depending on number of total barcode combinations used*). DNA undergoes a gap-fill reaction via Taq polymerase, in-vitro transcription via T7 RNA polymerase. RNA is purified and reverse transcribed using a random hexamer primer, then primed for second strand synthesis using primer complementary to the MED transposon. ME-B-only Tn5 was used to simultaneously fragment and adapter-tag 3’-end of cDNA to prepare for PCR indexing. (B) **Resulting library fragments** contain an r5 index added during targeted pA-Tn5 tagmentation and an i5 and i7 index added during PCR to enable retroactive demultiplexing of single cell reads.

### Multiplexing multiple antigens and single cell experimental conditions

Since combinatorial indexing begins with the addition of Index 1 to cells in a 96 well plate, multiple sample types (cells/antibody) can easily be multiplexed at this step then demultiplexed in-silico according to which Index 1 was added to the corresponding wells. As proof-of principle for our multiplexing scheme, we treated 5 separate pools of mouse F121-9 cells for antibodies against the histone post-translational modifications H3K27me3, H3K27ac, H3K9me3, serine-2 phosphorylated RNA pol II (Pol2S2p) and CTCF. Reads were aligned to the reference genome using bowtie2 (Langmead and Salzberg, 2012) and reads with low mapping-quality scores and PCR/optical duplicates were removed. Overall, we obtained data for 3740 single cells with a median of 93,180 mapped reads per cell (mapping rates >92%) and a mean of 92% unique (non-duplicate) reads, remarkably high-quality library complexity compared to existing methods (**Figure 4A**). These high alignment ratios, high quality of mapped reads, and excellent library complexity reflect the vastly improved read coverage resulting in exceptional quality of sequencing data obtained for the single cells. After filtering out cells with fewer than 1000 reads, we obtained 2770 cells with an average of 64,200 reads/cell (**Figure 4B-C**) that were used for further analysis.

**Figure 4.**
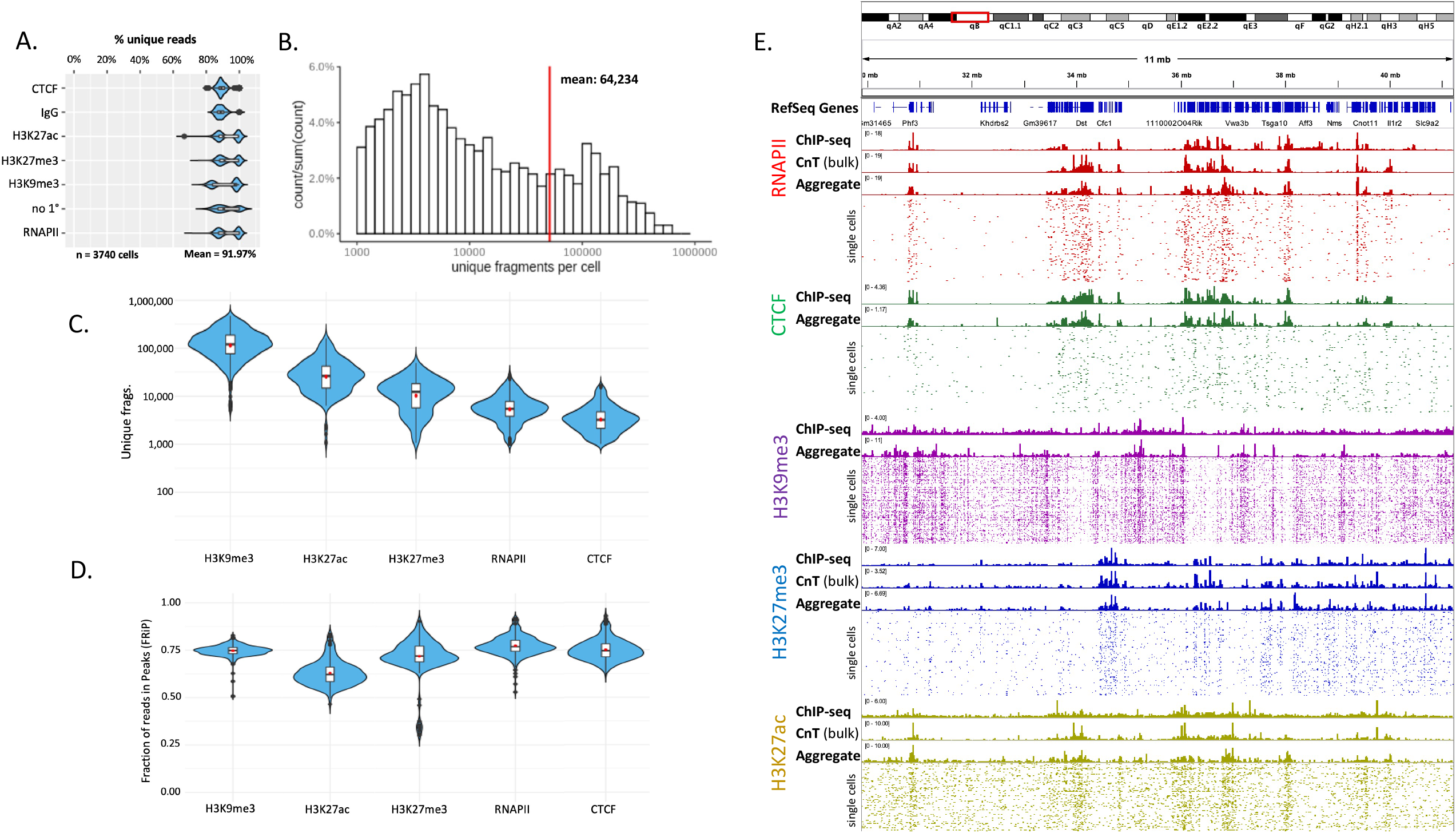
Multiplexed sciTIP-seq in mouse F121-9 ESCs. (A) Violin plots showing the percentage of unique (non-duplicated) reads per sample for each demultiplexed single-cell; (mean 92%). (B) Distribution of unique reads of 2,770 single cells in mouse ESCs after removal of cells with less than 1000 reads. Red line indicates the mean (64,234) unique fragments per cell. (C) Violin plots showing the number of unique reads per cell for each sample. (D) Violin plots showing the fraction of reads in single F121-9 cells falling within called peaks (FRiP) for each respective sample. (E) IGV track view across a 11Mb segment of the mouse genome for sciTIP-seq. Tracks show 100 single cells together with both pseudo-bulk (aggregate) and Bulk TIP/CnT/ChIP data (when available) at the representative loci for each sample – RNAPII, CTCF, H3K9me3 H3K27me3, H3K27ac. Bulk ChIP data are from ENCODE and 4D Nucleome consortium.

We visually inspected how the single-cell data agreed with bulk profiles by first creating a ‘pseudo-bulk’ sample by aggregating the reads from all the individual cells. The pseudo-bulk displayed similar profiling to those generated in Bulk TIP/ChIP/CnT (when available) samples and the reads from individual cells overwhelmingly fell within enriched regions of Bulk samples (**Figure 4E**). We then assessed signal-to-noise ratios of individual cells by pooling all reads from individual cells (per sample) into an aggregate sample and called peaks using MACS2 (Zhang et al., 2008) and measured the fraction of unique fragments per cell that fell within the called peaks (FRiP). Median FRiP scores of all single cells was 75% (**Figure 4D**), indicative of high signal-to-noise ratios equal-to or better than the best published methods.

### Single-chromosome mapping enabled by high coverage sciTIP data

In the vast majority of cases, single cell data is the average of two chromosome. The only way to overcome this obstacle is to either use haploid cells, for which there are very few cell lines, or to parse the single nucleotide polymorphisms (SNPs) that exist between maternal and paternal genomes in cases where they have been haplotype phased. Single-chromosome mapping of diploid cells requires much greater read depth to overcome the loss of reads that do not overlap SNPs. The extremely high read coverage of TIP-seq suggested that we might be able to map protein-binding sites on single chromosomes. As proof-of-principle, we parsed the data from F121-9, which harbor SNPs at a density of one per ∼150bp on average, that represent the polymorphisms between sub-species Mus musculus (129) and Mus castaneus (Cas). To parse alleles in RNAPII sciTIP data from hybrid F121-9 mESCs, we filtered for reads containing SNPs using a published algorithm (SNPsplit; https://www.bioinformatics.babraham.ac.uk/projects/SNPsplit/). Reads were aligned to their respective phased genomes and from there they were processed as in non-parsed sciTIP. Indeed, sites with substantial differences in occupancy between the Cas and 129 alleles were easily identified by visual inspection, and confirmed by comparing to parsed bulk CUT&Tag data (**Figure 5**).

**Figure 5.**
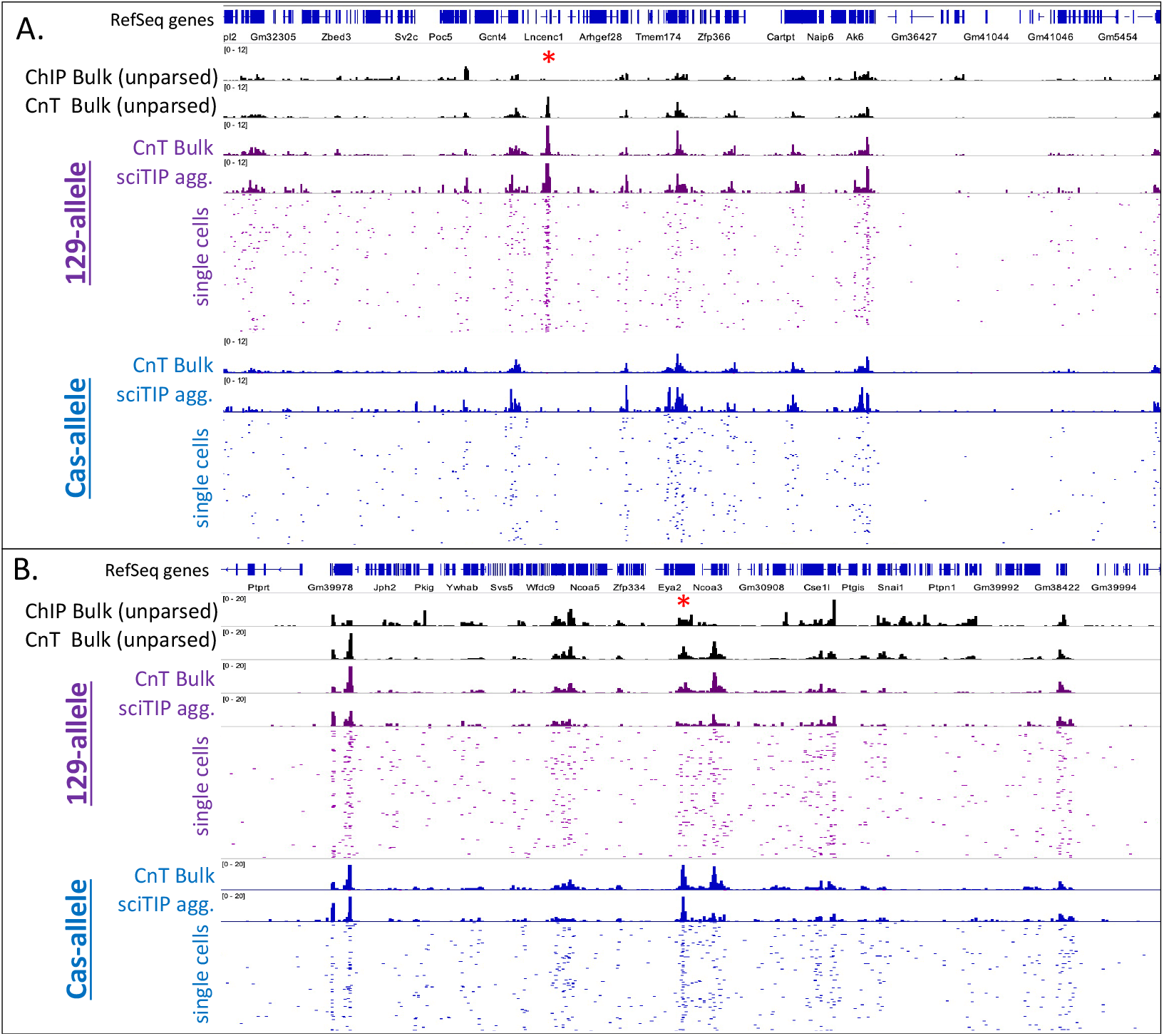
Single-chromosome mapping of RNAPII to parsed alleles in hybrid F121-9 mESCs. (A) IGV track view across a 9Mb segment of the mouse chromosome 13 comparing unparsed and parsed bulk CUT&Tag to parsed sciTIP aggregate and single cells. Tracks show unparsed bulk-ChIP (ENCODE), unparsed bulk-CUT&Tag, parsed bulk-CUT&Tag, parsed sciTIP pseudo-bulk (aggregate), and 100 parsed single cells. Red asterisks mark loci with differential RNAPII binding. (B) IGV track view across a 7Mb segment of the mouse chromosome 2. Everything else is the same as in panel B.

## DISCUSSION

Here, we show that TIP-seq produces single-cell sequencing data for both histones and TFs with dramatically higher read coverage per cell, substantially higher mappability and library complexity, and considerably lower background, compared to all current chromatin mapping methods. Tip-seq resembles CUT&Tag in that it uses Tn5 fused to protein A to insert a barcode adjacent to the protein binding site, but it differs in that the Tn5 MEDS inserted DNA contains a T7 promoter, and the insertion sites are linearly amplified by in vitro transcription with T7 polymerase, followed by cDNA library preparation prior to PCR indexing. Other single-cell methods traditionally require merging the reads of hundreds of single cells to acquire an aggregate profile that resembles bulk profiles. Remarkably, TIP-seq yields single-cell profiles resembling those of bulk samples with just a single sufficiently sequenced cell due to the high read coverage per cell.

A recent study (bioRxiv: Wu et al.) adapted single-cell CUT&Tag to use the droplet-based 10x genomics single-cell ATAC-seq platform. While it enables easy and automated single-cell partitioning and barcoding, it still requires access to specialized equipment and expensive commercial kits. They also reported a much lower throughput of only 2794 cells (1331 after filtering) (28% or 13%) of the expected 10,000 cells (as advertised by 10x Genomics). Nonetheless, they obtained CUT&Tag data for H3K27me3 in 1331 single cells that yielded a median of 1721 unique fragments/cell, whereas sciTIP-seq yielded 30-fold higher coverage at >52,300 unique fragments/cell for H3K27me3. A second study (bioRxiv: Bartosovic et al.) also adapted scCnT for the 10x Genomics platform, this time for H3K4me3, H3K27ac, H3K36me3 (active gene bodies), H3K27m3, and for transcription factors Olig2 and cohesion complex component Rad21. Again, library coverage was 10-fold lower than Tip-seq with a median <450 unique fragments for histone modifications. For transcription factors, they obtained a median of 48 and 240 unique reads per cell, as compared to Tip-seq yielding 4000 unique fragments/cell for transcription factors (100-fold and 10-fold higher, respectively).

Another recently developed method reported by 3 groups (ACT-seq, Carter et al., 2019; CoBATCH, Wang et al., 2019; Paired-Tag, Zhu et al., 2021) adapts CUT&Tag for combinatorial indexing. Unlike sciTIP-seq, these methods used PCR rather than IVT amplification. Carter et. al. reported ∼2500 unique reads/cell for 1246 cells with a read-mapping rate of 83% and only 15% were unique reads (duplication rate of 85%). sciTIP-seq yields 92% unique read rate and thus 25-fold more unique read/cell. Wang et. al. reported between 7500-12,000 unique reads/cell for 2161 cells (albeit cells with <3000 reads were removed from that statistic) and a duplication rate of 48%. Zhu et a. reported similar numbers of unique reads per cell (Table I; comparing Tip-seq to Paired Tag). By contrast, sciTIP-seq yields approximately 10-fold more unique reads/cell, and a far greater proportion of unique (non-duplicated) reads per library (92%) indicating that these libraries were not even sequenced to saturation/full potential (Table I; comparing Tip-seq to Paired Tag).

**Table I.** Library complexity: Unique reads/ cell comparison

|  | **<u>Paired-Tag</u>** | **<u>sciTIP-seq</u>** | **<u>Fold increase</u>** |
| --- | --- | --- | --- |
| **K27ac** | 4500 | 49,000 | 10.9x |
| **K27me3** | 2500 | 18,750 | 7.5x |
| **K9m3** | 17,500 | 121,000 | 6.9x |

In summary, TIP-seq offers high-quality (mapping rates), low background (FRiP) single cell libraries with substantially lower duplication rates (% unique reads), and thus a substantially improved per-cell library complexity compared to other single-cell chromatin mapping datasets (87-fold 10x Genomics scCnT; 25-fold ACT-seq; ∼10-fold CoBATCH and Paired-Tag). This unambiguously demonstrates the benefit of linear amplification. Moreover, the low duplication rates in our hands suggests these sciTIP-seq samples were not even sequenced to saturation thus, deeper sequencing may further improve the coverage of Tip-seq single cell data. It is likely that TIP-seq can achieve the read coverage necessary to perform studies of single cell variation in allele specific epigenomic features, such as both imprinted and random mono-allelic expression. Finally, TIP-seq will also be adaptable for other platforms, such as 10X genomics and ICELL8, and can be adapted to capture HiC, or RNA-seq and sciTIP protein mapping data from the same single cells.

## Acknowledgements

We want to thank K. Ahmad for early guidance and valuable comments on the manuscript, H. Kimura and T. Handa for training with ChIL-seq that inspired the design and implementation of TIP-seq, J. Turner for cell culture assistance, T Sasaki for assistance with sequence library pooling and handling, S. Miller and A. Brown for assistance with MiSeq pilot runs and Nextera Indexes, Y. Yang for help with Illumina NovaSeq6000 and bcl2fastq troubleshooting. This work was funded by National Institutes of Health grant R21 HG010403.

## Data Availability

Raw data is deposited in GEO XXXXXXXXX (**pending**) and data analysis scripts are available at https://github.com/dbartlett/sciTIP-seq2021

## Author Contributions

DAB and DMG conceived the experiments, DAB designed and executed all experiments and analysis, DAB and DMG wrote the manuscript, VD wrote cell demultiplexing scripts and SH provided pA-Tn5 and valuable consultation.

## Supplementary Tables

Supplemental table 1. List of oligos used in this study.

Supplemental table 2. NovaSeq 6000 RunInfo.xml and sample sheet.

## References

van Bakel, H., van Werven, F.J., Radonjic, M., Brok, M.O., van Leenen, D., Holstege, F.C.P., and Timmers, H.T.M. (2008). Improved genome-wide localization by ChIP-chip using double-round T7 RNA polymerase-based amplification. Nucleic Acids Res. 36, 21.

Baranello, L., Kouzine, F., Sanford, S., and Levens, D. (2016). ChIP bias as a function of cross-linking time. Chromosom. Res.

Brind’Amour, J., Liu, S., Hudson, M., Chen, C., Karimi, M.M., and Lorincz, M.C. (2015). An ultra-low-input native ChIP-seq protocol for genome-wide profiling of rare cell populations. Nat. Commun. 6, 6033.

Cao, J., Packer, J.S., Ramani, V., Cusanovich, D.A., Huynh, C., Daza, R., Qiu, X., Lee, C., Furlan, S.N., Steemers, F.J., et al. Comprehensive single-cell transcriptional profiling of a multicellular organism.

Cao, Z., Chen, C., He, B., Tan, K., and Lu, C. (2015). A microfluidic device for epigenomic profiling using 100 cells. Nat. Methods.

Carter, B., Ku, W.L., Kang, J.Y., Hu, G., Perrie, J., Tang, Q., and Zhao, K. (2019). Mapping histone modifications in low cell number and single cells using antibody-guided chromatin tagmentation (ACT-seq). Nat. Commun. 10.

Carter, B., Ku, W.L., Kang, J.Y., Hu, G., Perrie, J., Tang, Q., and Zhao, K. (2020). Author Correction: Mapping histone modifications in low cell number and single cells using antibody-guided chromatin tagmentation (ACT-seq) (Nature Communications, (2019), 10, 1, (3747), 10.1038/s41467-019-11559-1). Nat. Commun. 11, 1–1.

Chen, C., Xing, D., Tan, L., Li, H., Zhou, G., Huang, L., and Xie, X.S. (2017). Single-cell whole-genome analyses by Linear Amplification via Transposon Insertion (LIANTI). Science (80-.). 356, 189–194.

Cusanovich, D.A., Daza, R., Adey, A., Pliner, H.A., Christiansen, L., Gunderson, K.L., Steemers, F.J., Trapnell, C., and Shendure, J. (2015). Multiplex single-cell profiling of chromatin accessibility by combinatorial cellular indexing. Science (80-.).

Eberwine, J., Yeh, H., Miyashiro, K., Cao, Y., Nair, S., Finnell, R., Zettel, M., and Coleman, P. (1992). Analysis of gene expression in single live neurons. Proc. Natl. Acad. Sci. 89, 3010–3014.

van Galen, P., Viny, A.D., Ram, O., Ryan, R.J.H., Cotton, M.J., Donohue, L., Sievers, C., Drier, Y., Liau, B.B., Gillespie, S.M., et al. (2016). A Multiplexed System for Quantitative Comparisons of Chromatin Landscapes. Mol. Cell 61, 170–180.

Grosselin, K., Durand, A., Marsolier, J., Poitou, A., Marangoni, E., Nemati, F., Dahmani, A., Lameiras, S., Reyal, F., Frenoy, O., et al. (2019). High-throughput single-cell ChIP-seq identifies heterogeneity of chromatin states in breast cancer. Nat. Genet. 51, 1060– 1066.

Hainer, S.J., Bo Skovi C, A., Mccannell, K.N., Rando, O.J., and Fazzio, T.G. (2019). Profiling of Pluripotency Factors in Single Cells and Early Embryos. Cell 177, 1319– 1329.

Harada, A., Maehara, K., Handa, T., Arimura, Y., Nogami, J., Hayashi-Takanaka, Y., Shirahige, K., Kurumizaka, H., Kimura, H., and Ohkawa, Y. (2019). A chromatin integration labelling method enables epigenomic profiling with lower input. Nat. Cell Biol.

Hashimshony, T., Wagner, F., Sher, N., and Yanai, I. (2012). CEL-Seq: Single-Cell RNA-Seq by Multiplexed Linear Amplification. Cell Rep. 2, 666–673.

Hoeijmakers, W.A.M., Bártfai, R., Françoijs, K.J., and Stunnenberg, H.G. (2011). Linear amplification for deep sequencing. Nat. Protoc. 6, 1026–1036.

Kaya-Okur, H.S., Wu, S.J., Codomo, C.A., Pledger, E.S., Bryson, T.D., Henikoff, J.G., Ahmad, K., and Henikoff, S. (2019). CUT&Tag for efficient epigenomic profiling of small samples and single cells. Nat. Commun. 10, 1930.

Ku, W.L., Nakamura, K., Gao, W., Cui, K., Hu, G., Tang, Q., Ni, B., and Zhao, K. (2019). Single-cell chromatin immunocleavage sequencing (scChIC-seq) to profile histone modification. Nat. Methods 16, 323–325.

Lake, B.B., Chen, S., Sos, B.C., Fan, J., Kaeser, G.E., Yung, Y.C., Duong, T.E., Gao, D., Chun, J., Kharchenko, P. V., et al. (2018). Integrative single-cell analysis of transcriptional and epigenetic states in the human adult brain. Nat. Biotechnol. 36, 70– 80.

Langmead, B., and Salzberg, S.L. (2012). Fast gapped-read alignment with Bowtie 2. Nat. Methods 9, 357–359.

Lareau, C.A., Duarte, F.M., Chew, J.G., Kartha, V.K., Burkett, Z.D., Kohlway, A.S., Pokholok, D., Aryee, M.J., Steemers, F.J., Lebofsky, R., et al. (2019). Droplet-based combinatorial indexing for massive-scale single-cell chromatin accessibility. Nat. Biotechnol. 37, 916–924.

Marinov, G.K. (2018). A decade of ChIP-seq. Brief. Funct. Genomics 17, 77–79.

Minkina, A., and Shendure, J. (2019). Expanding the single-cell genomics toolkit. Nat. Genet. 51, 931–932.

Mulqueen, R.M., Pokholok, D., Norberg, S.J., Torkenczy, K.A., Fields, A.J., Sun, D., Sinnamon, J.R., Shendure, J., Trapnell, C., O’Roak, B.J., et al. (2018). Highly scalable generation of DNA methylation profiles in single cells. Nat. Biotechnol.

Ramani, V., Deng, X., Qiu, R., Gunderson, K.L., Steemers, F.J., Disteche, C.M., Noble, W.S., Duan, Z., and Shendure, J. (2017). Massively multiplex single-cell Hi-C. Nat. Methods.

Rooijers, K., Markodimitraki, C.M., Rang, F.J., De Vries, S.S., Chialastri, A., De Luca, K.L., Mooijman, D., Dey, S.S., and Kind, J. Simultaneous quantification of protein-DNA contacts and transcriptomes in single cells.

Skene, P.J., Henikoff, J.G., and Henikoff, S. (2018). Targeted in situ genome-wide profiling with high efficiency for low cell numbers. Nat. Protoc. 13, 1006–1019.

Sos, B.C., Fung, H.L., Gao, D.R., Osothprarop, T.F., Kia, A., He, M.M., and Zhang, K. (2016). Characterization of chromatin accessibility with a transposome hypersensitive sites sequencing (THS-seq) assay. Genome Biol.

Tang, G.Q., Bandwar, R.P., and Patel, S.S. (2005). Extended upstream A-T sequence increases T7 promoter strength. J. Biol. Chem. 280, 40707–40713.

Vitak, S.A., Torkenczy, K.A., Rosenkrantz, J.L., Fields, A.J., Christiansen, L., Wong, M.H., Carbone, L., Steemers, F.J., and Adey, A. (2017). Sequencing thousands of single-cell genomes with combinatorial indexing. Nat. Methods.

Wang, Q., Xiong, H., Ai, S., Yu, X., Liu, Y., Zhang, J., and He, A. (2019). CoBATCH for High-Throughput Single-Cell Epigenomic Profiling. Mol. Cell.

Weiner, A., Lara-Astiaso, D., Krupalnik, V., Gafni, O., David, E., Winter, D.R., Hanna, J.H., and Amit, I. (2016). Co-ChIP enables genome-wide mapping of histone mark co-occurrence at single-molecule resolution. Nat. Biotechnol.

Zhang, B., Zheng, H., Huang, B., Li, W., Xiang, Y., Peng, X., Ming, J., Wu, X., Zhang, Y., Xu, Q., et al. (2016). Allelic reprogramming of the histone modification H3K4me3 in early mammalian development. Nature 537, 553–557.

Zhang, Y., Liu, T., Meyer, C.A., Eeckhoute, J., Johnson, D.S., Bernstein, B.E., Nussbaum, C., Myers, R.M., Brown, M., Li, W., et al. (2008). Model-based analysis of ChIP-Seq (MACS). Genome Biol. 9.

Zhu, C., Zhang, Y., Li, Y.E., Lucero, J., Behrens, M.M., and Ren, B. (2021). Joint profiling of histone modifications and transcriptome in single cells from mouse brain. Nat. Methods 1–10.

## BioRxiv Referenes

Wu, S.J., Furlan, S.N., Mihalas, A.B., Kaya-Okur, H., Feroze, A.H., Emerson, S.N., Zheng, Y., Carson, K., Cimino, P.J., Keene, C.D., Holland, E.C., Sarthy, J.F., Gottardo, R., Ahmad, K., Henikoff, S., and Patel, A.P. (2020). Single - cell analysis of chromatin silencing programs in developmental and tumor progression. bioRxiv doi: https://doi.org/10.1101/2020.09.04.282418

Bartosovic, M., Kabbe, M., and Castelo-Branco, G. (2020). Single-cell profiling of histone modifications in the mouse brain. bioRxiv doi: https://doi.org/10.1101/2020.09.02.279703

